# Reassessing taxonomy and virulence in the *Fusobacterium nucleatum* group – Rebuttal of *Fusobacterium animalis* clades “*Fna* C1” and “*Fna* C2”, genome announcement for *Fusobacterium watanabei* and description of *Fusobacterium paranimalis* sp. nov

**DOI:** 10.1101/2025.03.20.644344

**Authors:** Audun Sivertsen, Diego Forni, Cristian Molteni, Joanna Bivand, Grete Dimmen, Manuela Sironi, Øyvind Kommedal

## Abstract

There is a considerable interest in the association between *Fusobacterium animalis* and colorectal cancer (CRC). Recently, it was suggested that this association is valid only for a distinct clade of *F. animalis* (*Fna* C2) and that *F. animalis* strains belonging to another clade (*Fna* C1) are only associated with the oral cavity. It was further suggested that this made *Fna* C1 a natural comparator when looking for candidate genes associated with the pathogenicity of *Fna* C2. Based on such comparisons, three candidate operons enriched in CRC were suggested to explain the strong colorectal tumor association of *F. animalis*.

In the present paper we show that major taxonomic errors invalidate the existence of two distinct clades of *F. animalis* and that *Fna* C1 is simply a rediscovery and misclassification of *Fusobacterium watanabei*. We further reassess the phylogenetic structure of the entire *Fusobacterium nucleatum* group encompassing *F. animalis* and all known closely related species and confirm the current taxonomy using contemporary phylogenetic principles. We also describe a novel *Fusobacterium* species more closely related to *F. animalis* than any other known species, for which we propose the name *Fusobacterium paranimalis* sp. nov..

We further searched for the three proposed candidate virulence operons of *F. animalis* across the entire *F. nucleatum* group and show that some or all of these are present in all other species except *F. watanabei*. We also observe considerable variability of Type 5 secretion systems (T5SS) by subtype and abundance across the *F. nucleatum* group.

**Importance:** It is known that *Fusobacterium animalis* is able to survive within colorectal tumors. Recently, it was proposed that only one “clade” of *Fusobacterium animalis* could be found in colorectal tumors, and that another “clade” within the same species was instead only found in the oral cavity, and that differences in gene content could explain the habitat difference. We here show that these “clades” are two separate species by sequencing the type strain of the oral cavity-associated species, which is *Fusobacterium watanabei*. We also revisit other related species within the “*Fusobacterium nucleatum* group” to confirm that they are separate species, exemplified by presenting a new species, *F. paranimalis* sp.nov., which genetically is more related to *F. animalis* than any other known species including *F. watanabei*. We look at gene content of the entire group, and conclude that known virulence genes cannot fully explain *F. animalis* cancer association.

## Introduction

There is significant interest around the connection between *Fusobacterium animalis* and colorectal cancer (CRC). In a recent study comparing presumed *F. animalis* strains isolated from the human oral cavity and CRC tumors, Zepeda-Rivera *et al*. (2024) (1) conclude that *F. animalis* can be divided into two distinct clades, called “*Fna* C1” and “*Fna* C2”, whereof only “*Fna* C2”, which includes the *F. animalis* type strain, is associated with CRC. By categorizing “*Fna* C1” as a clade of *F. animalis* not associated with CRC they frame it as the natural comparative taxonomic group for studying the pathogenic properties of the cancer associated *“Fna* C2*”*. The study further implies that the standard species-level of bacterial identification in diagnostic microbiology is inadequate for research on the role of the microbiome in cancer and has generated substantial attention.

We found several flaws and taxonomic misconceptions invalidating the tale of two clades within the *F. animalis* species: (i) when classifying “*Fna* C1” isolates as a lineage of *F. animalis*, the authors fail to adhere to basic taxonomic principles even though these principles are correctly described in the article text; (ii) the authors do not discuss that “*Fna* C1” is at least as closely related to *Fusobacterium vincentii* as it is to *F. animalis* although this is clear from ANI calculations provided in their Supplementary Table S4; and (iii) they have not included all relevant *Fusobacterium* species in their analyses, thereby failing to acknowledge that “*Fna* C1” is simply a rediscovery and misclassification of the valid species *Fusobacterium watanabei*.

*Fusobacterium watanabei*, most closely related to *F. vincentii* and *F. animalis*, was announced by Tomida *at al.* in 2021 (2) and accepted as a validly published species outside of the International Journal of Systematic and Evolutionary Microbiology later the same year (3). Although Tomida *et al*. report ANIb and dDDH values for its closest neighboring species within the *Fusobacterium nucleatum* group below 92.2% and 49.5% respectively, no genome was published to accompany the novel species announcement. At present, only partial sequences of the *16S rRNA* gene (1400 bp), together with partial sequences of *rpoB* (700 bp), *gyrB* (942 bp), and a zinc protease gene (551 bp) is present in nucleotide databases to support the taxonomy. As the type-strain is not represented in genome databases such as NCBI RefSeq or GTDB (4), later genome references for other strains of *F. watanabei* lack the correct species designation. In GTDB, *F. watanabei* is currently called *Fusobacterium nucleatum_J*.

While the failure of Zepeda-Rivera *et al*. to acknowledge the existence of *F. watanabei* might be explained by the lack of an available whole genome reference, their rationale for classifying it as a clade of *F. animalis* instead of a new species remains an enigma. The authors have been challenged on these matters (5, 6) but so far no correction has been published for their paper (7). We therefore acquired the type-strain of *F. watanabei* from the CCUG strain collection (*F. watanabei* CCUG 74246T) and sequenced it using a combination of Illumina and Oxford Nanopore sequencing. We then re-analyzed the *Fusobacterium* genomes from the study by Zepeda-Rivera *et al*. together with the novel genome for *F. watanabei* CCUG 74246T using contemporary phylogenetic tools and principles. We also included the genome of a *Fusobacterium* strain (Vestland19) isolated from blood culture, recently mentioned as a putative new species in a study by Bivand et al (8). Finally, we reassessed whether the CRC-enriched fusobacterial operons described by Zepeda-Rivera *et al*. are unique to *F. animalis*, or are also present among other species of the *F. nucleatum* group.

## Results and discussion

### Re-assessment of the *Fusobacterium nucleatum* group taxonomy

The “*Fusobacterium nucleatum* group” is a term encompassing *Fusobacterium animalis*, *Fusobacterium nucleatum*, *Fusobacterium polymorphum*, *Fusobacterium vincentii* (i.e. the four former subspecies of *Fusobacterium nucleatum* sensu lato), and other closely related species (9). It is a relevant term since the *16S rRNA* gene, commonly used for identification in many settings including diagnostic microbiology, discriminates poorly between several of these species. To evaluate the *F. nucleatum* group genetic diversity, all genomes used by Zepeda-Rivera *et al.* were downloaded. In addition, we sequenced the type strain of *F. watanabei*, CCUG 74246T, along with a putative novel *Fusobacterium* species isolated from a positive blood culture at the Dept. of Microbiology, Haukeland University Hospital. Finally, we included available genomes of *F. hwasookii* (N=8) and *F. simiae* (N=4), two additional species closely related to *F. animalis* also not included in the data analyses in the Zepeda-Rivera *et al*. paper. All genomes were included in an all-vs-all ANI calculation using SKANI. A heatmap highlighting the canonical 95% ANI species threshold (Fig. 1A) shows that CCUG 742426T clusters together with the strains erroneously suggested to compose *F. animalis* Clade 1 (“*Fna* C1”), confirming that these strains are indeed *F. watanabei*. A closer look at ANI differences between CCUG 74246T and all other strains in this dataset (Fig. 1B) reiterate findings from Zepeda-Rivera *et. al.* that *F. watanabei* and *F. animalis* share ∼93% ANI between the groups, supporting that they are distinct species as per the 95% ANI definition. Fig. 1B also shows that *F. vincentii*, *F. animalis* and *F. watanabei* strains are all within ∼93% ANI from each other, bringing into question why only *F. animalis* and *F. watanabei* were selected as comparators in the virulence experiments conducted, and why *F. vincentii* was considered a distinct species and not *F. watanabei (*aka “*Fna* C1”). The ANI-analyses also support that “*Fusobacterium* strain Vestland19” represents a novel *Fusobacterium* species more closely related to *F. animalis* than any other known species in the *F. nucleatum* group. For this species we therefore propose the name *Fusobacterium paranimalis* sp. nov.

**Figure 1.**
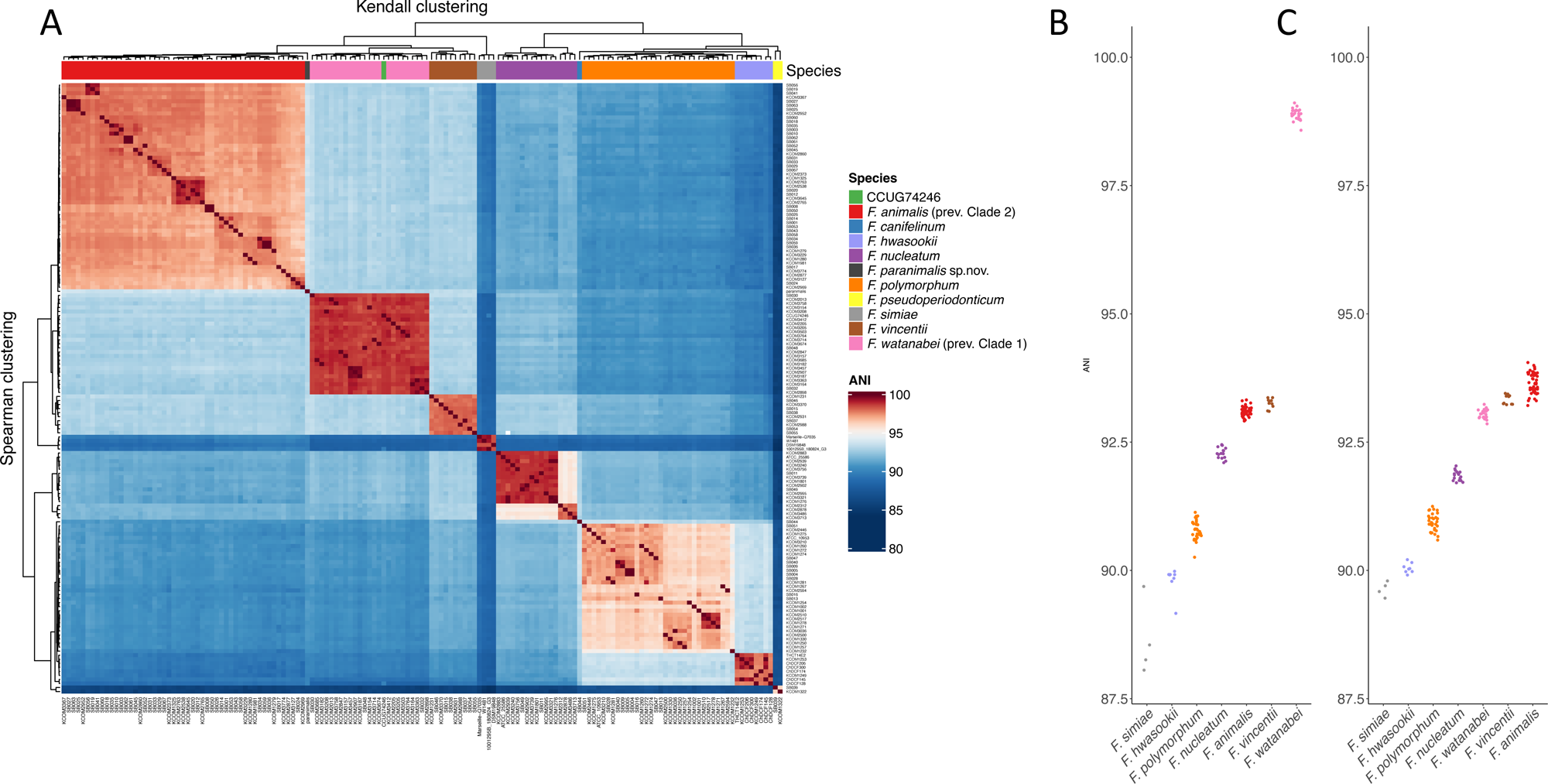
ANI comparisons between *Fusobacterium* spp. (a) Heatmap of ANI differences between genomes within the *Fusobacterium nucleatum* group as determined by SKANI, with color shading highlighting the canonical 95% ANI species delineation rule. *F. watanabei* type strain CCUG 74246T is highlighted. (b) Jitter plot showing ANI of species with multiple strains to CCUG 74246T. (c) Jitter plot showing ANI of species with multiple strains to type strain of *F. paranimalis* sp. nov.

### Beyond ANI - the Biological Species Concept (BSC) and the *Fusobacterium nucleatum* group

Although ANI analysis is widely used to define bacterial species, this approach is based on arbitrary thresholds. As a consequence, alternative strategies, based on the Biological Species Concept (BSC), have been proposed (10–12). The BSC defines a species as a group of interbreeding individuals that remain reproductively isolated from other groups. In the case of bacteria, gene-flow discontinuities can be used to delineate biological species (10, 13). Because the interruption of gene flow is only partially dependent on the degree of genetic divergence (14), ANI-defined species do not necessarily correspond to species defined by BSC. We thus applied two methods based on BSC to delineate the composition of biological species within the *F. nucleatum* group.

We first used the whole dataset of *Fusobacterium* genomes as an input for PopCOGenT (populations as clusters of gene transfer) (10), which can detect recent gene-flow discontinuities that delineate species. PopCOGenT compares the length distribution of identical regions between pairs of genomes to that expected under a model of clonal evolution, and it provides a measure referred to as “length bias”. PopCOGenT identified 12 genetically isolated ecological units and revealed clusters with excellent congruence to ANI-defined species, with no detectable gene-flow among them (Fig. 2A). PopCOGenT results thus support that *F. watanabei* and *F. animalis* are distinct biological species and that *F. paranimalis* represents a novel species. These data also indicate that, in accordance with ANI analysis, the *F. nucleatum* genomes belong to two distinct ecological units, as is the case for the two *F. pseudoperiodonticum* genomes. The genetic heterogeneity of *F. pseudoperiodonticum* has previously been noticed (6).

**Figure 2.**
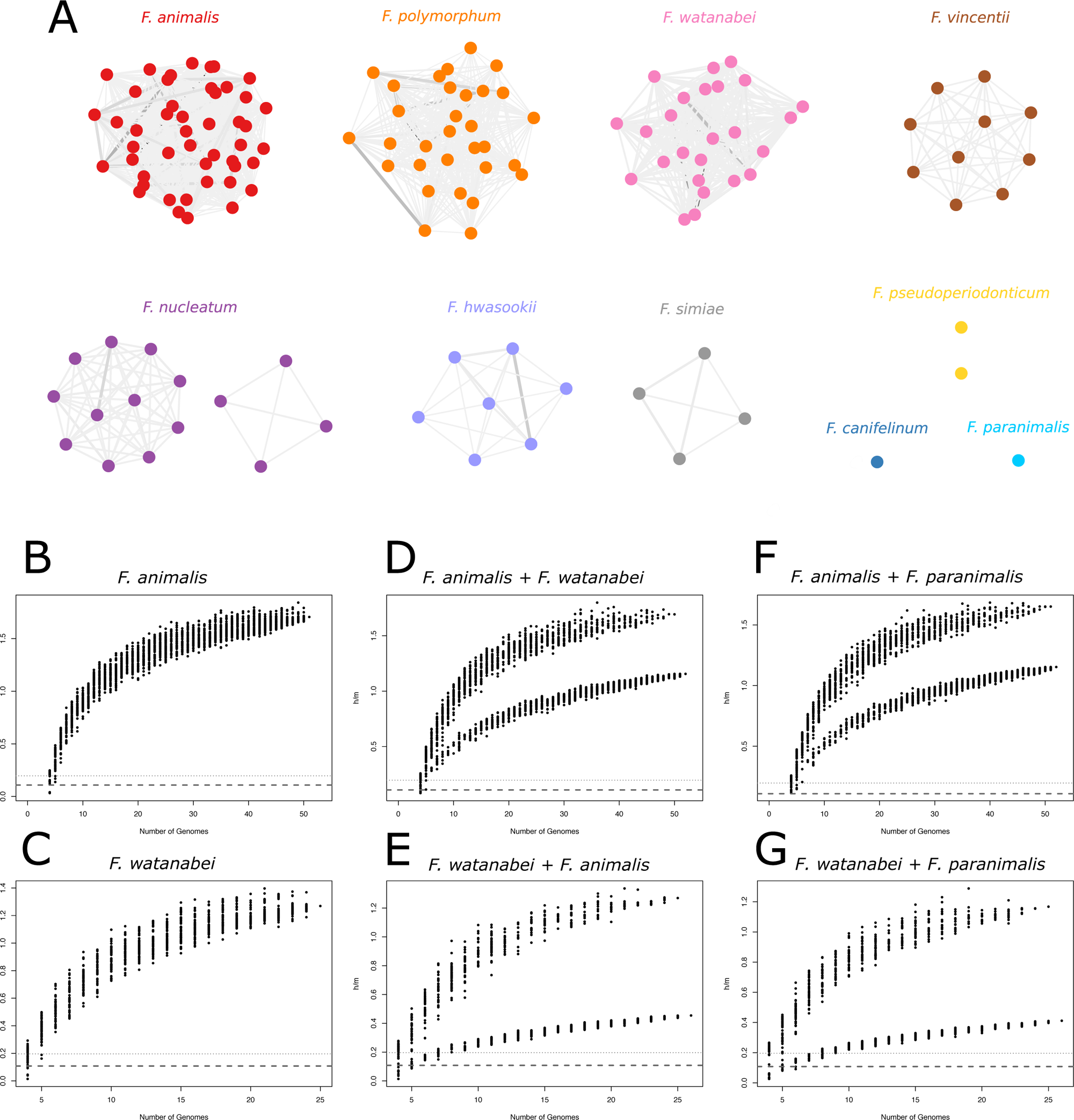
Gene flow between *Fusobacterium* species. (a) Gene flow network of *Fusobacterium* genomes. Nodes represent bacterial strains and edges represent the inferred amount of gene flow between them (expressed in terms of length bias). Edge width and color are proportional to the amount of gene flow between pairs of genomes. Clonal clusters (i.e. strains too closely related) were collapsed in single nodes. (b) Homologous recombination between *Fusobacterium* species. Homoplasy/mutation ratios of different *Fusobacterium* species are plotted. A single curve indicates the presence of one species, whereas the presence of a second lower curve indicates that two different species are present in the sample. The *F. watanabei* type strain and a random *F. animalis* strain were used as test lineage in the *F. animalis* + *F. watanabei* and in the *F. watanabei* + *F. animalis* comparisons, respectively.

We next wished to corroborate these results using an approach that relies on the detection of homoplasies, which can persist for long times and thus measure the effect of both recent and historical gene-flow (13–15). Specifically, we applied ConSpeciFix (13), which measures the ratio of homoplastic (h) to non-homoplastic (m) polymorphisms across core genomes. Based on the h/m parameter, the program estimates whether the tested genomes form a coherent unit, that is a biological species. We focused on *F. animalis* and *F. watanabei*, which showed high h/m values, indicative of widespread gene-flow among core genomes within each species (Fig. 2B, C). The inclusion of *F. watanabei* CCUG 74246T to the *F. animalis* population or the addition of one *F. animalis* genome to the *F. watanabei* population resulted in sharp drops of h/m values, confirming that *F. animalis* and *F. watanabei* are distinct biological species (Fig. 2D, E). Similarly, inclusion of the *F. paranimalis* genome to either species caused a sharp decrease in the h/m ratio, confirming that *F. paranimalis* is genetically isolated from *F. animalis* and *F. watanabei* (Fig. 2F, G).

We next aimed to determine whether methods based on core genome alignments were suitable to reconstruct genetic relationships within the *Fusobacterium nucleatum* complex. We thus used the Genome Taxonomy Database Toolkit (GTDB-Tk) (16) to extract the sequences of 120 core genes present in these fusobacterial genomes. The alignment of the core genes was used to generate a neighbor-net split network, which recapitulated the phylotaxonomic order determined by the analyses above (Fig. S1). Thus, *F. animalis* and *F. watanabei* were clearly separated, and *F. paranimalis* clustered closer to *F. animalis*, in line with ANI analysis. The split of *F. nucleatum* into two lineages was also observed in the network. Similar results to those described above were obtained when we used the Gubbins (Genealogies Unbiased By recomBinations In Nucleotide Sequences) program to construct a phylogenetic tree that accounts for the effect of recombination (Fig. 3) (17), where all mentioned groups clustered independently of each other.

**Figure 3.**
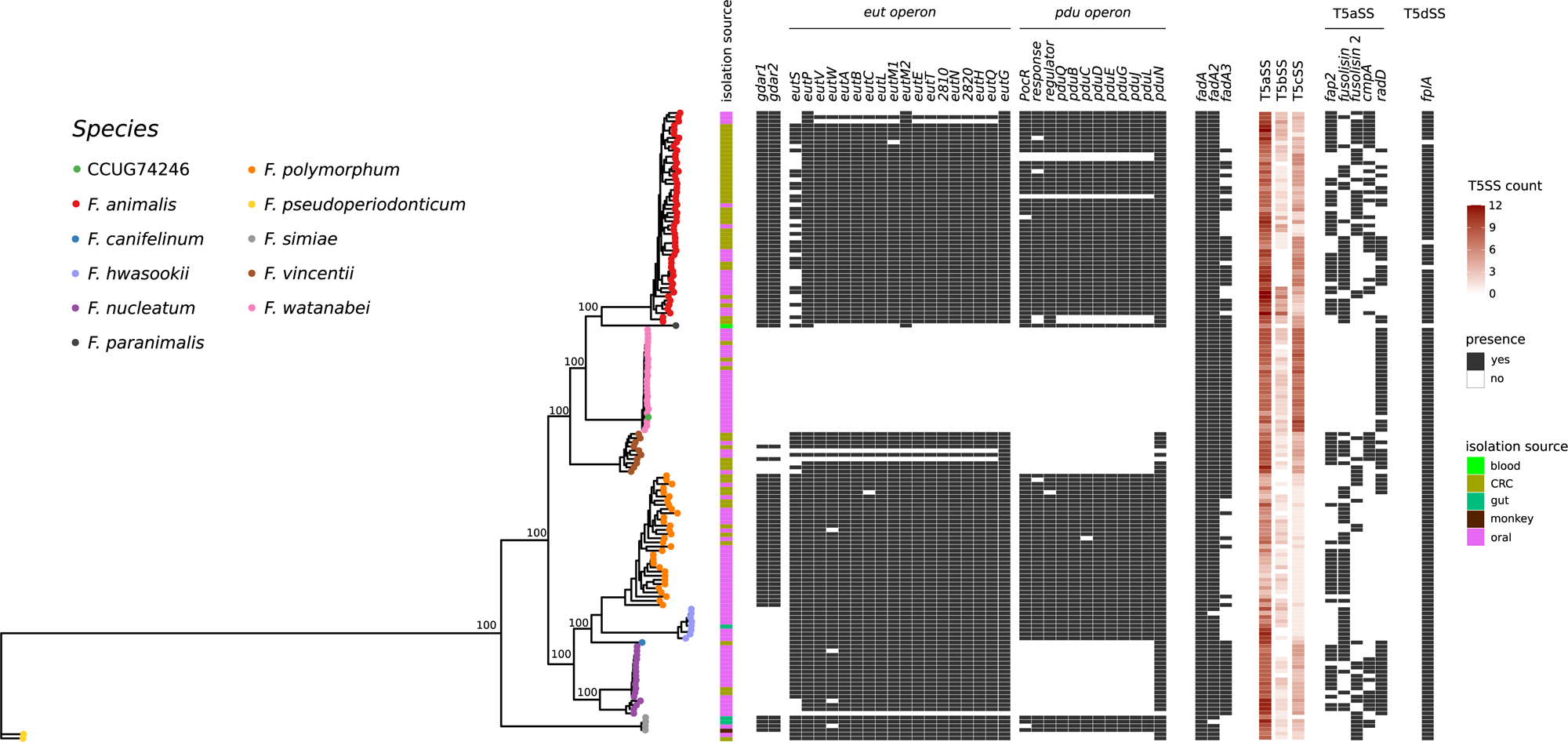
Distribution of virulence factors among *Fusobacterium* species. A recombination-aware phylogenetic tree of core genomes calculated by GUBBINS is shown, color-coded by species. Bootstrap support values (expressed in percentage) for relevant nodes are reported. A gene presence/absence matrix for the virulence factors calculated by PPanGGOLiN is also reported, along with information about strain isolation sources and the counts of Type V secretion system components estimated by macsyfinder. Color categories are explained in the included legends.

A recent work by Connolly et al (2025) (18), which included all genomes in Zepeda-Rivera *et al.* (2024) but left out *F. simiae* and *F. hwasooki*, in phylogenetic and PopCOGenT analyses found identical species distributions as we did. They did not name *Fna* C1 as *F. watanabei*, but several observations like the one presented by Connolly *et al.* highlights that species designations up until this point have been inconsistent and likely inaccurate.

### Re-assessment of virulence factor presence in all *Fusobacterium nucleatum* group members

Zepeda-Rivera *et al*. framed their “*Fna* C1” (*F. watanabei*) as a natural comparator for the CRC associated “*Fna* C2” (*F. animalis*) and do not report whether other closely related species harbour their suggested virulence operons *eut*, *pdu* and the GDAR acid resistance system. Using their rationale, *F. paranimalis* sp. nov. should represent an even more suitable comparator species in power of being even more closely related to “*Fna* C2” (*F. animalis*). However, candidate virulence operons should not be put forward based on a comparison of only two taxons within a larger group of closely related species. Also, occasional presence of other *Fusobacterium* spp. (like *F. nucleatum, F. vincentii, F. polymorphum, and F. pseudoperiodonticum)* in CRC samples (19, 20) could indicate commonalities in virulence gene content among these species.

We therefore searched for the three candidate virulence operons from *F. animalis* in all other closely related species by running PPanGGOLIN on all strains. Briefly, the software clusters proteins into families of likely orthologs based on provided identity and coverage scores, and keeps track of co-localisation of each protein within the genomes included in the analysis. These protein families were then annotated using representative protein sequences of stated candidate virulence factors, and presented as a presence/absence matrix along with the Gubbins tree (Fig. 3).

Interestingly, *F. polymorphum* and *F. simiae* contain all three operons along with *F. animalis*. These two species are not associated with CRC, although *F. polymorphum* appears associated with oral dysplasias (21). These species should therefore perform similarly to *F. animalis* in biochemical tests performed by Zepeda-Rivera *et al*.. Further, the *F. hwasooki* strains lack only the GDAR system, whereas *F. nucleatum* sensu stricto, *F. pseudoperiodonticum,* and *F. vincentii* lack both GDAR and *pdu* operons but harbor the *eut* operon. The single genome of *F. paranimalis* sp. nov. contains the *pdu* and GDAR operon, but lacks *eut*.

*F. watanabei* is the only species lacking all the candidate virulence operons. Our finding that these operons are also harboured by other *Fusobacterium* species, makes us conclude that their presence alone cannot answer why only *F. animalis* is so strongly associated with CRC. The results together imply that there still are other important pieces to the puzzle, and that although including more species in pan genome comparisons may complicate the picture, it might also unravel novel genes and operons important to *Fusobacterium* pathogenicity with higher precision.

The gene(s) encoding secreted FadA (22), appearing in three separate forms (FadA1-3), is present in all strains of our collection and therefore is a likely ubiquitous virulence factor of the *F. nucleatum* group. FadA is shown to be secreted from the cell by T5aSS/autotransporter Fap2 (22), but, as we and others (21, 23, 24) have found, the T5aSS/autotransporters in Fusobacterium spp. are not easily classified into discrete groups. PPanGGOLIN reported several gene families within the “autotransporter” group, and numerous and genetically different genes are annotated as “autotransporter domain-containing protein” making it difficult to disentangle orthology/paralogy relationships. Single protein searches using e.g. Diamond allowed single reference T5SS proteins (Aim1, RadD, Fap2, fusolisin, FplA) to map to several loci within each strain, inflating the count, but also limiting the potential count of T5SS due to arbitrary cutoffs to a limited set of reference proteins which may not represent the full repertoire of T5SS within Fusobacterium spp. For instance, the prototypic Fap2 protein is only found in some strains within the F. animalis population (Fig. 3), which speculatively indicates that also other T5aSS have involvement in FadA transport across the cell wall.

We therefore used Macsyfinder v.2 (25) to identify the number of T5SS in each strain using TXSScan, which identifies secretion systems through Hidden Markov Model (HMM) protein profiles. In general, a wide variability in the number of T5SS components was observed, both among and within species (Fig. 3). Moreover, an interesting pattern emerged where T5aSS proteins seem enriched in *F. animalis, F. vincentii, F. hwasooki, and F. nucleatum* sensu stricto compared to other species. *F. watanabei* is instead enriched in T5cSS relative to other species, another feature distinguishing it from *F. animalis*.

A striking difference is seen between T5SS abundance and type distributions, where only one T5dSS (*FplA*) (26) seems ubiquitous within the *F. nucleatum* group without large variations. In contrast, T5aSS are more present in numbers, but each “type”, here clustered by moderate cut-offs of 70% sequence identity and 80% coverage, is relatively sparsely present between the strains and species we analysed.

Speculatively, incompatibility dynamics may be at play, exemplified by the two PPanGGOLIN protein families being most similar to fusolisin (27); these two groups shared ∼55% protein sequence identity and could be found in all analysed species except *F. watanabei*. If one of the fusolisin types was present, the other type was seldom concurrently seen. These two proteins were the most similar within the N-terminal and C-terminal ends, and most variation within the substrate-binding sequence normally located in the middle (28). Determining whether the two fusolisin protein families share the same substrate, like histones (29), falls outside the scope of this article.

In sum, host-microbe interactions between *Fusobacterium* spp. and human tissues do not simply hinge on one or few virulence factors. An important task is to first identify the important niches and ecological correlates of each species within the group, before detangling what accessory genes do in each *Fusobacterium* species, in each context. In this regard, Connolly *et al*. (2025) (18) again highlight that *F. animalis* and *F. polymorphum* seem enriched in cancer tissues and gut lesions in

Crohn’s disease by analysing metagenomes from such sites, and that *F. vincentii* together with *F. polymorphum* are associated with gingival plaque. *F. animalis, F. vincentii and F. polymorphum* were also enriched in stools from patients with type two diabetes. Additionally, interestingly, as we did, Connolly *et al.* identified a bifurcation in the *F. nucleatum* sensu stricto population, of which the smallest group was found more often in stool samples than the larger group.

### Genome announcement for Fusobacterium watanabei CCUG 74246T

*Fusobacterium watanabei* CCUG 74246T was *de novo* assembled using Oxford Nanopore (ONT) combined with Illumina NGS data. The GC content was 27% and the resulting contig of 1981216 bp included 47 tRNAs, 15 rRNAs, one CRISPR array and 1816 CDSs when assembled with Bakta. As seen in Fig. 1A and B, CCUG 74246T clusters with other *F. watanabei* strains, and all other *F. watanabei* members share ∼>98% ANI with CCUG 74246T.

### Description of *Fusobacterium paranimalis* Vestland19T

*Fusobacterium paranimalis* sp. nov. (Gr. prep. *para*, beside, near, like; L. gen. n. *animalis*, of an animal; *paranimalis* resembling (*Fusobacterium*) *animalis*) is a long slender fusiform anaerobic Gram-negative rod (Fig. S2), closely related to *F. animalis*. It grows with greyish colonies (0.5-1 mm) after 48 hours of incubation in a strict anaerobe atmosphere at 35 degrees Celsius. The *16S rRNA* gene shares 99.8-100% identity with multiple uncultured references from the human oral cavity in GenBank including “*Fusobacterium sp.* oral taxon C10 clone DD027” (Accession GU429671) submitted by the Human Oral Microbiome Project (www.homd.org). It is likely a commensal species from the human oral microbiota.

The type strain is *F. paranimalis* Vestland19T (Strain numbers in DSMZ and NCTC to be revealed later). The strain was isolated in a single blood culture bottle from a patient with a suspected but unconfirmed bacterial infection in 2019 who recovered without antimicrobial treatment. Most likely, the finding represented a transient bacteremia originating from the oral cavity. Using MaldiTOF-MS (Bruker MALDI Biotyper database MBT Compass Library 2023 (12438MSP)) for identification gives a low confidence identification with score 1.8 for *F. nucleatum*. The draft genome size is 2.4 MB, encoding 2289 genes. The G+C content is 26,9 mol% .

## Conclusions

A meaningful search for candidate virulence operons that can explain a perceived species-specific pathogenicity necessitates a correct and comprehensive phylogenetic description coupled with the inclusion of all relevant taxonomic groups in the investigation. Our results confirm the current taxonomy for the *F. nucleatum* group, including *F. watanabei*. We decisively reject the division of *F. animalis* into two distinct clades as suggested by Zepeda-Rivera *et al.*, a suggestion that was never supported by their data. This further dismantles their argument for using “*Fna* C1” (*F. watanabei*) as the natural comparator for “*Fna* C2” (*F. animalis*) when searching for candidate genes of potential importance in *F. animalis* CRC tumor invasion. Our expanded analyses show that the candidate operons set forward by Zepeda-Rivera *et al*. are not unique to *F. animalis* and consequently cannot solely explain the enrichment of *F. animalis* in colorectal tumors. Finally, we confirm the discovery of a novel species within the *F. nucleatum* group more closely related to *F. animalis* than any other known member of the cluster for which we propose the name *F. paranimalis*.

## Methods

### Sequencing of strains *F. watanabei* CCUG 74246T and *F. paranimalis* sp. nov. Vestland19T

The type strain CCUG 74246T was purchased from CCUG in 2024. *F. paranimalis* sp.nov. Vestland19 was retrieved from a blood culture bottle in the clinical laboratory. After storage in −80°C, the strains were cultivated on acumedia® blood agar plates (Neogen). Genomic DNA extraction was performed from bacterial colonies using the ELITe Ingenius DNA extractor (ELITechGroup, Turin, Italy) for Illumina sequencing and TANBead Maelstrom 4800 (Taiwan Advanced Nanotech, Taoyuan, Taiwan), for Oxford Nanopore (ONT) sequencing. The ONT library protocol was Rapid Sequencing Kit V14 (Oxford Nanopore, UK), sequencing was performed using a R10.4.1 flongle. Reads were trimmed and basecalled using Dorado v.7.4.14 invoked through MinKNOW. Illumina reads were also produced, using the Nextera XT DNA Library Preparation Kit (Illumina) and the Illumina MiSeq System. The genome was subsequently *de novo* assembled using Hybracter v0.10.1 (30) hybrid-single with short-read polishing of the flye assembly. Long-read coverage was downsampled to 100x by hybracter, short-read coverage was ∼91 for CCUG 74246T and ∼224 for Vestland19. The genomes were annotated using Bakta v. 1.9.2 (31).

### ANI comparisons

The largest collection of *F. watanabei* with other *Fusobacterium* spp. isolates was found in the bioproject associated with the article of Zepeda-Rivera *et al.* 2024; PRJNA549513. Using Skani v.0.2.2 (32), all strains with >∼85% ANI similarity to the type strain CCUG 74246T were included in the ANI pair-to-pair comparison to create Fig.1A in R using ComplexHeatmap (33). The species within the *F. nucleatum* group which fell within ∼90% ANI and contained >4 strains were subsampled to create the dotplot in Fig. 1B and C using ggplot2 (34).

### Core gene analyses

All genomes in our dataset were used as input data for GTDB-Tk, v2.1.1 (16). This tool allows the identification and extraction of sequence information for a set of 120 single copy marker genes that are conserved among bacteria using whole genome sequences as input data. A concatenated alignment based on these core genes was then generated using MAFFT v7.475 (35) with default parameters. A neighbor-net split network was built using SplitsTree v4.16.2 (36) based on HKY85 distances, using parsimony-informative sites, and removing gap sites.

The core gene alignment was also screened for the presence of recombination signals using Gubbins (v.3.0.0) (17). Polymorphic sites outside those recombination regions were then extracted and used to build a recombination-aware phylogeny using fasttree (37) for generating the starting tree and IQ-TREE for the final tree (38). The best substitution model was automatically selected by the ModelFinder tool implemented in IQ-TREE. Robustness of the final tree was assessed using 1000 bootstrap replicates. All other parameters were set as default.

### BSC-based methods

We applied two different methods based on the BSC: PopCOGenT (10) and ConSpeciFix (13).

PopCOGenT defines species boundaries based on recent horizontal gene transfer by searching for stretches of high sequence identity between any pair of genomes. Thus, this tool detects gene-flow across the entire genomic space, including both core and accessory genes. PopCOGenT estimates “length bias”, a measure of the observed length distribution of identical regions between pairs of genomes compared to a null expectation of non-recombinogenic evolution (10). Based on this measure, PopCOGenT infers population predictions and generates networks of gene flow, with strains as nodes and the length bias as a measure of their relationship. We ran PopCOGenT using all the 151 *Fusobacterium* strains in our dataset and we plotted gene flow networks using Cytoscape v3.9.1 (39).

ConSpeciFix instead focuses on a core set of genes shared by a group of genomes. In particular, this tool defines which genes are core, then evaluates the ratio of homoplastic/recombinant polymorphisms (h) to mutations (m) across them. Finally, to determine species membership, a genome sequence is compared to a set of genomes that are already defined as a single species and the h/m ratio is calculated for an increasing larger number of genomes. If the tested strain belongs to a different species, h/m ratios will decrease (13).

To understand the relationship between *F. animalis*, *F. watanabei*, and the proposed *F. paranimalis* sp. nov., we ran ConSpeciFix using the “personal comparison” method. We ran two preliminary analyses using *F. animalis* or *F. watanabei* separately, and then each species was analysed adding one strain of the other. Then, we analysed these two populations by adding the *F. paranimalis* genome.

### Identification of virulence factors

The pangenome of all Fusobacterium strains was calculated by running PPanGGOLIN v2.2.1 (40), using as input protein sequences and genBank annotation files. Gene family clustering was then computed through the MMseqs2 tool implemented in PPanGGOLIN and with thresholds set to 70% for sequence identity and 80% for sequence coverage, without performing the defragmentation step. All other parameters were set as default. Single gene searches for T5aSS (Fad2, Aim1, RadD, Fusolisin, FplA) were done by extracting representative protein sequences from strain ATCC 23726 and searching each genome using diamond blastx (41) with 70% coverage and 70% identity to reference.

The generated pangenome was further used to create a gene presence/absence matrix of virulence genes, selected on the basis of Zepeda-Rivera et al. (1) and by information from the genBank annotation files. Finally, the presence/absence matrix was plotted along with the Gubbins tree using the ggtree R package.

Identification of all the best candidate Type V secretion system components (i.e. the monomeric autotransporters (T5aSS), the two-partner secretion system (T5bSS), and the trimeric secretion system (T5cSS)) was conducted with MacSyFinder (v2.1.4) (42) by running the TXSScan macsy-models (25) and selecting proteomes as “unordered”.

## Data availability

CCUG 74246T and *F. paranimalis* sp.nov. are available in bioproject PRJEB85514. CCUG 74246T has accession GCA_965119615.1 and chromosome accession OZ221964; Illumina reads are available as SRA experiment ERX13642389, ONT reads are available as SRA experiment ERX13777046. For Vestland19, the complete chromosome accession is to be specified later. Separate accession numbers for the complete 16S rRNA and *rpoB* genes will also be specified later.

## Acknowledgments

The work was supported by the Italian Ministry of Health -“Ricerca Corrente” program (to DF and CM).

---

**Supplementary Figure S1.**
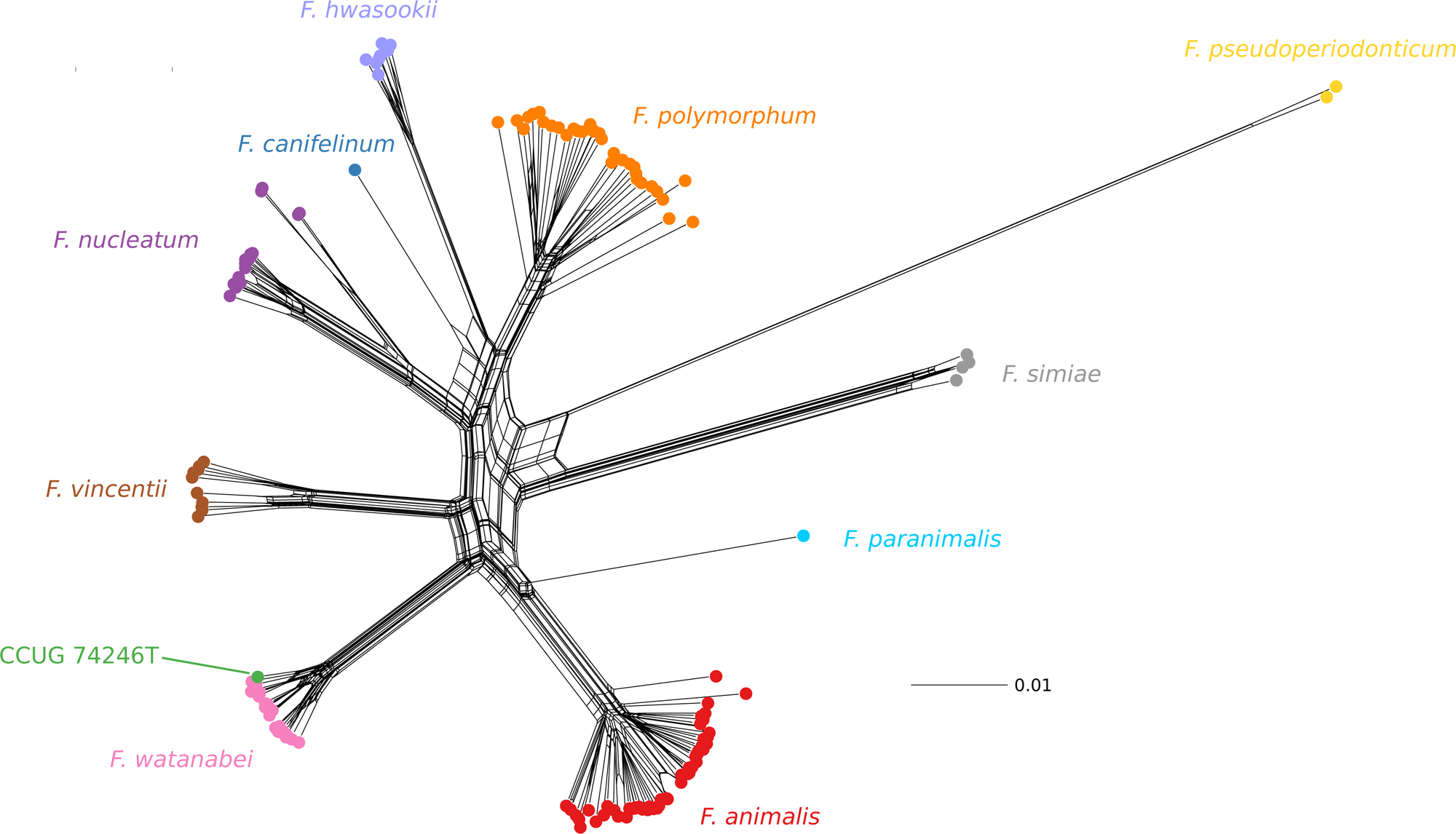
Neighbor-net split network of Fusobacterial core genomes. Each sequence is shown as a dot, color-coded by species. Within the *F. watanabei* cluster, the placement of the type strain CCUG 74246T is shown.

**Supplementary Figure S2.**
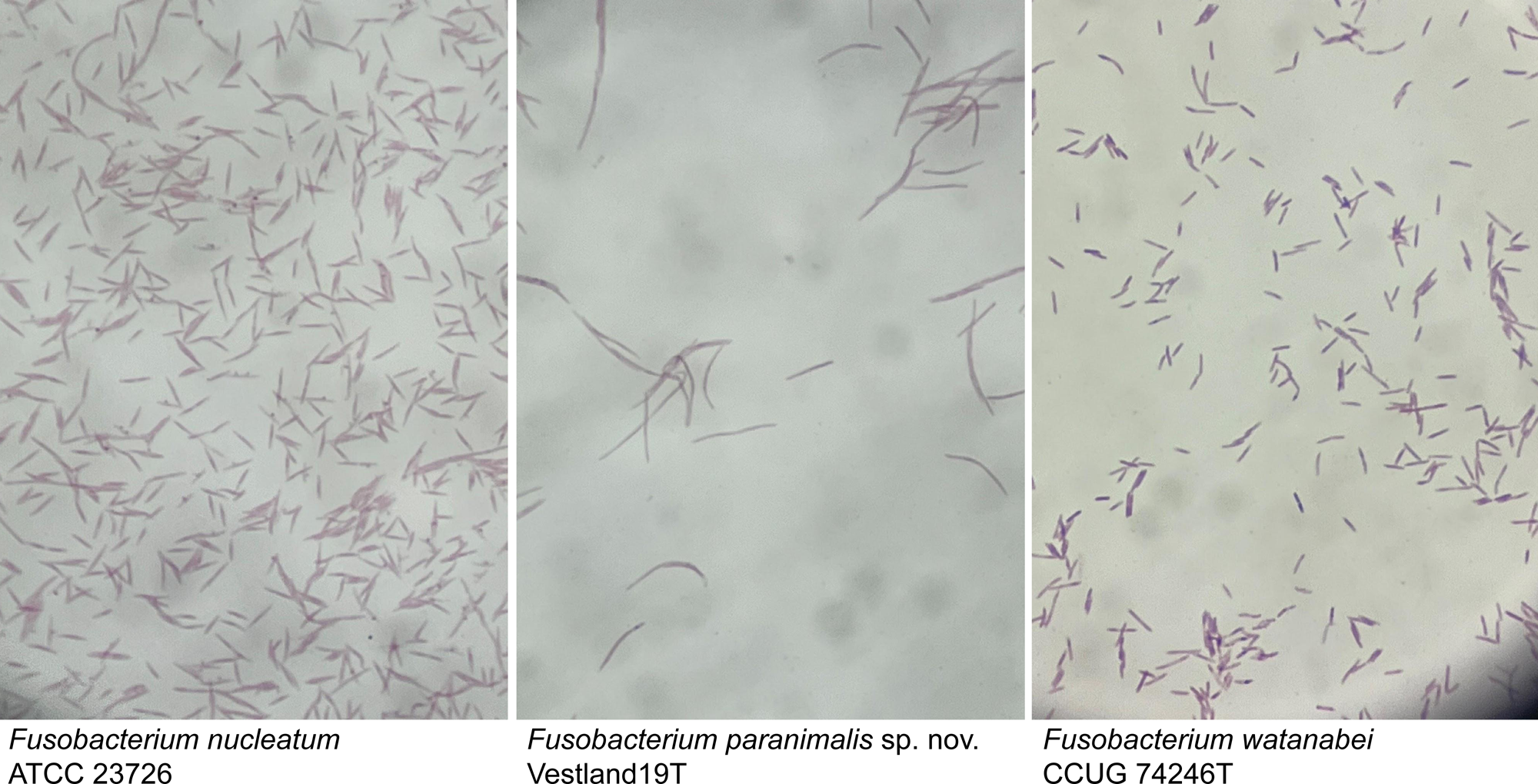
Light microscope morphology of *F. nucleatum, F. paranimalis,* and *F. watanabei*.

